# Bacterial invasion of the pancreas revealed after analyses of the pancreatic cyst fluids

**DOI:** 10.1101/064550

**Authors:** Vilvapathy Narayanan, Wesley K. Utomo, Marco J. Bruno, Maikel P. Peppelenbosch, Sergey R. Konstantinov

## Abstract

The involvement of bacterial translocation (BT) in the promotion of carcinogenesis has gained a considerable attention in the last years. At this point however BT has not been studied in the context of pancreatic cystic lesions and their development into pancreatic ductal adenocarcinoma. The aim of the study was to analyze if bacteria are present in pancreatic cyst fluid (PCF) collected from patients with suspected pancreatic cysts. Total DNA was isolated from sixty nine PCF. The occurrence of bacteria in PCF was analyzed using bacterial 16S rRNA gene-specific PCR-based method followed by sequence identification and quantitative PCR assay tuned up to different pathogenic and commensal human bacteria. Forty-seven out of sixty-nine samples (68%) were found positive for harboring bacterial 16S rRNA gene. Follow up sequencing analyses of the PCR products revealed that bacterial species related to *Fusobacterium spp., Bacteroides spp.,* and *Bacillus spp.* were predominating the PCF samples. The results suggest that specific bacteria can translocate to the pancreas and become detectable in the PCF.

## Introduction

The incidence of pancreatic cystic lesions (PCL) in the general population is 2.4 percent. Neoplastic pancreatic cysts represent a risk for developing pancreatic ductal adenocarcinoma ^1^ (PDAC) and account for up to 5% of the total incidence of pancreatic cancerous lesions ^2,3^. The clinical challenge in PCL is to identify signs of progressive neoplastic transformation in order to perform a surgical resection before a malignancy develops.

The vast majority of cysts nowadays are asymptomatic and coincidental finding at cross sectional imaging done for other reasons than cyst related symptoms.The percentage of cystic lesions being resected is increasing with the clinical intention of preventing the development of PDAC. Different types of cystic lesions are presently characterized by Grutzmann *et. al,* and Farrell *et. al.,* which include the intra-ductal papillary mucinous neoplasm (IPMNs), mucinous cystic neoplasms (MCN), serous cyst adenomas (SCA), and pseudocysts ^4,5^. IPMN’s and MCN’s pose a risk of developing into carcinoma of which IPMNs are being more prevalent compared with MCN’s ^2,6^. IPMNs are further classified as main branch, side branch or mixed types, based on the extent of involvement of the pancreatic ductal system ^5^. Presently, there are no validated biomarkers to identify cystic lesions that require surgical resection. Although only up to three percent of the PCL patients would develop cystic lesions to malignancy, ten percent of the PCL patients are resected ^5^ suggesting the need for more superior clinical tests and better patients’ stratification prior to surgery.

Currently, the decision for resection of PCL and/or continued monitoring are made according to the Sendai guidelines after evaluation of different clinical tests ^7-9^. The available clinical tests include different biochemical analysis, cytology, pathological identification, endoscopic ultra sonography (EUS), and radiological diagnosis such as endoscopic retrograde cholangiopancreatography (ERCP), magnetic resonance cholangiopancreatography (MRCP) and whole body computerized tomography (CT). The inter-observer agreement, however, within and between different modalities remains moderate. Therefore, a set of preoperative biochemical analyses have been increasingly used in clinical decision making. This includes the study of cyst fluids and serum for the characteristic presence of carcinoembryonic antigen (CEA), cancer antigen 19.9 (CA-19.9), tumour associated glycoprotein 72-4 (CA-72-4), cancer antigen 15-3 (CA-15-3), pancreatic amylase, mucin antigens and along with cysts characteristics ^10,11^. Other PCL tests are based on specific analyses of different genetic modalities like K-RAS mutation and miRNA, but are still under investigation awaiting further clinical validation ^2,6,12,13^. Though different diagnostic tests could help in decision making, none of them is sensitive and specific enough to make the right decisions for all patients involved. Additional markers are therefore are urgently needed to improve the management of PCL.

The human gut microbiome has emerged recently as an important environmental factor linked to the development of different intestinal and extra-intestinal malignancies ^14-16^. Several members of the intestinal microbiome have been implicated in the bacterial translocation (BT) that could occur during different diseases of the pancreas ^17-21^. Numerous studies suggest that BT takes place via the mesenteric lymph nodes route, followed by hepatic portal route, and trans mural or biliary or duodenopancreatic reflux ^18^. This may initiate and/or accelerates intestinal leakage with a subsequent lowering of host immune response followed by dissemination of commensal gut microbiota and their by-products to other organs leading to sepsis and major multiple organ failures ^21,22^. BT has been demonstrated in induced acute pancreatitis in mice and other animal models where BT takes place immediately towards the pancreatic duct causing necrosis in the exocrine system of pancreas ^19,23^. Most of the BT has been described to originate from the small intestine rather than from the colon ^19^. BT-associated intestinal pathobionts include different strains belonging to *Escherichia coli, Enterobacter cloacae, Enterobacter fecalis, Proteus mirabilis,* and *Pseudomonas* species that have been involved in the necrotizing and acute pancreatitis ^18^. Although there is ample evidence that BT takes place in various pancreatic diseases, not much has been studied nor understood about the bacterial presence in pancreatic cyst and/or cyst fluids. In this discovery phase study we have attempted to establish the bacterial occurrence in the pancreatic cystic lesions.

## Materials and Methods

**Patient samples and Pancreatic cyst fluid collection.** A cohort of 103 patients with suspected cystic lesions was established between the period of 2008-2013 of which sixty nine samples were randomly selected for this discovery phase study (Table 1). The pancreatic cyst fluids (PCF) were collected after a signed informed consent from these patients who were undergoing endoscopic ultra sound fine needle aspiration (EUS-FNA) at the department of Gastroenterology & Hepatology, Erasmus MC, The Netherlands. The PCF were collected, immediately transferred to the lab and stored at -150oC.

**DNA isolation.** Approximately 300 μl from sixty nine PCF samples were used for total DNA isolation. After bead beating (Fast Prep®-24 Instrument) the supernatant and pellet were separated by centrifugation at 13 000 rpm for 1 min and used for total DNA isolation using the Wizard DNA isolation kit as specified by the manufacturer’s protocol (Catalogue no. A1620, Promega BNL B.V, The Netherlands). Isolated DNA was equilibrated in the DNA rehydration solution from the kit and quantified on nanodrop-2000 spectrophotometer (Isogen Life Science BV, De Meern, The Netherlands). PCF DNA was diluted to 1ng/μl for the PCR analyses and subsequently stored at -20°C.

**(q)PCR analyses.** Total DNA isolated from the PCFs were used for bacterial 16S RNA gene detection using both conventional PCR and qPCR. For the conventional PCR based method GoTaq^®^ Flexi DNA polymerase kit was used (Promega BNL B.V, The Netherlands). qPCR was performed on IQ5 machine (Biorad, Bio-Rad Laboratories, Inc. Hercules, CA, United States of America) using Syber Green amplification kit (SYBR Select master mix, CFX, ABI, Life technologies, the Netherlands). All primers are listed in Table 2. All the primers were analysed and confirmed for the specificity to bacterial 16S rRNA and mismatch with the human mitochondrial 16S rRNA. 16S rRNA gene products from *Acinitobacteria spp., Anaerococcuss spp., Bacillus spp., Bacteroides spp., F. nucleatum, Propionibacterium spp.* and *Staphylococcus spp.* were used to generate PCR standards from 10 ng to 0.00001 ng concentrations thereby achieving total of seven standards. Water was used as negative control.

**16S RNA gene sequencing analyses.** Classical Sanger sequencing method was done in order to identify the bacterial 16S rRNA genes present in the PCF. The PCR products generated from universal primers of 16S rRNA 16S bacterial gene were sequenced and then after identification of specific bacteria which are pathogenic commensals and as well involved in cancer development were used for further sequencing. The *Fusobacterium spp., Bacteroides spp., Anaerococcus spp., Acinitobacteria spp., Propionibacterium spp., and Staphylococcus spp.* Specific 16S rRNA primers were used and sequenced (Table 2) through LGC Genomics GMBH, Germany.

**Fluorescent In-situ Hybridization.** The depraffinization of the slides were performed as the protocol used in our lab ^24^, once depraffinized the tissues slides are subjected to PBS washing for one minute. Modified proteinase K treatment was done under incubation at 45° C for 30 minutes to reduce the background staining (proteinase K − 20 μg/ml 1:10 dilution) (Sigma Aldrich, the Netherlands) ^25^. The slides are then dehydrated for 3 min in 50, 70, and finally 96% ethanol. 500 microliters of hybridization buffer (0.9 M NaCl, 20 mM Tris-HCl [pH 7.5] 0.1% [wt/vol] sodium dodecyl sulfate) were applied and incubated at 37° C for 30 minutes. The hybridization buffer is washed off in the prewarmed PBS at 37° C and new pre-warmed hybridization buffer is added with the FITC labelled probes containing 1 picomole of FITC-labeled 16S rRNA probes (EUB338 and *F. nucleatum)* was applied and incubated at 37° C for 18 hours. After hybridization, the slides were washed in 50 ml washing buffer (0.4X SSC buffer for 2 minutes followed with 2X SSC buffer for 1 minute, SSC buffer prepared as per the protocol by Zordan A et.al ^25^ and washed with PBS 1X for 1 minute and DAPI (1μg/ml) is added for counter staining of the nucleus and stained for exactly 3 minutes. The slides are washed with PBS 1X and air dried. All the staining are done in dark. The air dried slides are then examined using confocal microscope (LEICA, The Netherlands).

**Statistical methods.** All the statistical analysis were done using excel and Graphpad Prism 5.0.

The CT values in X axis were plotted against the serially diluted standards in Y axis to find the intercept Y using the exponential trend line for the CT values as per the excel formula. The R^2^ is also determined for the same plot. The Y intercept is used to calculate the ng of 16S rRNA for the unknown samples of *Fusobacterium spp., Bacteroides spp., Anaerococcus spp., Acinitobacteria spp., Propionibacterium spp.,* and *Staphylococcus spp..* Once the ng for each sample is obtained they are used for calculating absolute copy number of 16s rRNA using the following formula,

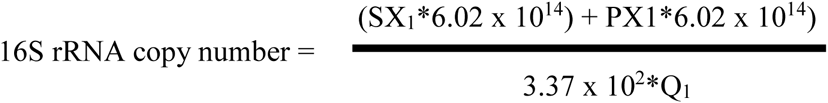

Where in SX_1_ and PX_1_ – S – stands for supernatant, P – stands for the pellet, X_1_ – stands for the sample 1, 6.02 x 10^14^ – nano gram to Daltons, 3.37 x 10^2^ molecular weight of one nucleotide in Daltons, Q1 – stands for the qPCR - product in number of nucleotides. The final conversion of 16S rRNA copy number to 1ml of PCF is calculated via 16S rRNA copy number multiplied by 3.33 (300 μl to 1ml) 300 μl of PCF fluid was used for total DNA isolation.

## Results

**Study sample characteristics.** Sixty nine PCF samples were used for this study (Table 1). PCF were characterized as different types of IPMN’s (39.1%), which included main branch IPMN (2.9%), mixed type IPMN’s (5.8%), side branch IPMN’s (17.4%) and multifocal side branch IPMN’s (5.8%), and unclassified IPMN’s (7.2%). Another types of PCF included in the study were mucinous cystic neoplasms (18.8%), pseudocysts (13.0%), serous cyst adenomas (13.0%). Finally, various other types of cystic lesions, gastrointestinal stromal tumour (GIST), neuroendocrine tumour (NET), and PCF without a definite clinical diagnosis were lumped into a separate category “other” PCF (15.9%) Table 1.

**Bacterial DNA presence in pancreatic cyst fluids.** We have found that 47 (68.11%) of the 69 samples were positive for bacterial DNA originating from different bacteria (Table 3). Various PCFs harboured different percentage of positive samples ranging from 69.2% of all MCN’s, followed by IPMN’s with 66.7% and pseudocysts (66.7%), SCA’s (55.6%), whereas in the group of others which included GIST, NET and clinically undefined samples were 71.4% predominantly positive for bacterial DNA (Table 6). This confirms the bacterial DNA present in the PCF’s.

**Bacterial DNA quantification in pancreatic cyst fluids.** We have used qPCR experiments to confirm the presence of typical intestinal bacteria using specific primer sets targeting *Bacteroides spp., Anaerococcus spp., Acinitobacterium spp., Propionibacterium spp.,* and selected pathobionts including *F. nucleatum, and Staphylococcus spp.* The copy number quantified for the total bacteria and the specific bacteria selected based on 16S rRNA qPCR were adjusted to ml of pancreatic cyst fluid are listed in Table 4. Based on the copy number the 16S rRNA were ranked the bacteria as represented in Table 5, which shows the pseudocysts (5.07 x 10^8^) shows the highest copy number for bacteria and the lowest also seen in SCA (3.53 x 10^5^), corroborating the presence of bacteria in pseudocysts as seen in earlier reports. The average bacterial 16S rRNA copy number /ml of PCF was 1.08 x 10^7^ with most occurring in pseudocysts (5.71 x 10^7^) followed with MCN (1.15 x 10^7^), IPMN (2.04 x 10^6^) others (GIST, NET and clinically undefined – 1.92 x 10^6^), and SCA (7.91 x 10^5^) (Figure 1). One sample in pseudocysts, showed an abnormal bacterial load of 5.07 x 10^8^ compared to other samples (Table 4). Similarly, there are change in the number of 16S rRNA copy number between various grades of dysplasia (graded according to the most atypical area in the lesion) in IPMN in the decreasing order: 2.46 x 10^6^ in no dysplasia, low grade dysplasia have 1.75 x 10^6^, Moderate dysplasia 9.67 x 10^5^ and carcinoma in situ 8.30 x 10^5^ (Fig 2, no statistical difference). qPCR on PCF with mucinous cystic neoplasms have also demonstrated a higher number of bacterial 16S rRNA gene in the samples where no dysplasia is present (Fig 3).

When the partial gene bacterial 16S rRNA (1464 nucleotide) normal PCR were conducted using universal primers (table-2) only 68.11% of the samples showed positive bacterial population presence. Whereas the quantification via qPCR of bacterial 16S rRNA (193 nucleotides) primers shown in table-2, shows presence of bacterial 16S rRNA in 100% of the samples.

**Sequencing of the bacterial 16S rRNA sequences.** PCR products generated using universal 16S rRNA, and primers specific for *F. nucleatum, Bacteriodes spp., Anerococcus spp., Acinitobacterium spp., Propionibacterium spp.,* and *Staphylococcus spp.* were subjected to sequencing to confirm the qPCR results (Table 6). The sequencing results have demonstrated that *F. nucleatum* is present in 13 out of all 69 PCF samples (18.84%). Other predominating bacterium involved in PCF was *Bacillus spp.* which was present in 16 out of 69 (23.19%) samples. The presence of other bacteria was also noted which included *Ruminococcus spp., Staphylococcus spp., Caldimonas spp., Arthrobacter spp., Acinetobacter spp., Bacteroides spp., Orpinomyces spp. Anaerococcus spp* (combined data from Tables 3 and 6).

**Fluorescent In-situ Hybridization (FISH).** FISH was used as an additional analysis to confirm the presence of bacteria in PCF. Selected slides were blindly analysed checked and scored for bacterial presence by two personal specialized in confocal microscopy. We have confirmed it by the confocal imaging of the FISH slides for the bacterial presence Figure 4. Importantly, our control samples, negative for bacterial presence in the PCF’s were also negative in the tissue slices generated after resection (data not shown). On the opposite side, the tissue samples from patients with PCF positive for bacterial DNA were also positive for bacterial presence in the tissue, the cysts borders and the duct borders (Figure 4).

## Discussion

In the current study we have demonstrated that 68 percent of the PCF harbour different types of bacteria. The possible BT into the pancreatic cysts involve also bacteria related to *F. nucleatum* implicated in adenoma to carcinoma transformation in epithelial cells ^26^. Although EUS-FNA is not a sterile procedure, 1/3 (32 percent) of the PCF samples were negative for bacterial DNA on our cultivation-independent bacteriological findings. Nonetheless EUS-FNA limitations for bacterial analyses need to be discussed. Before sampling cyst fluid, the needle tip is exposed to the content in the stomach or gut lumen before being punctured through the gut wall into the pancreatic cyst. The presence of bacterial 16S rRNA in all the 69 samples taken for this study, observed as in the qPCR is due to the above said reasons of EUS-FNA being an non-sterile procedure and also due to the bacterial products transferred via the other contaminants of the cyst fluids, like blood which is plausibly carrying the bacteria DNA via the bacteria ingested macrophages and other immune cells. To nullify and further account for that when interpreting our data, we have performed confocal imaging for bacteria confirming that the tissue samples resected from patients with PCF positive for bacterial DNA were also positive for bacterial presence in the tissue. Furthermore, many bacteria in the stomach like *H. pylori* and streptococci were not found in our PCF analyses supporting the notion that specific bacterial community may be associated with PCF.

BT involvement into the pancreatic abscess, necrosis, pancreatitis, and pancreatic cyst have also been reported mainly in isolated case studies, but never in large cohort analyses ^22,27-29^. Numerous case studies identified BT to pancreas were caused by the commensal bacteria and fungi. Pancreatic infections mainly arise from translocation of bacteria from the small bowl, and rarely from the colon and oropharyngeal route as demonstrated by study on *Veillonella* and *Bifidobacterium spp.* which were identified in pancreatic abscess ^28^. A study of Brook et al., has identified 158 bacterial species from pancreatic abscess of which 77 isolates were aerobic and rest 81 were anaerobic bacteria ^30^. Most commonly detected microorganisms in pancreatic pseudocysts include often not only opportunistic bacteria like *E. coli, Enterobacter spp., Klebsiella spp.,* and *Staphylococcus spp.,* but also fungal isolates including *Candida albicans* (15 case studies) ^27^. Importantly, EUS FNAB procedure caused serious *Clostridium perfringens* infections in 5 patients leading to pancreatitis and pancreatic cyst formation, which required surgical interventions ^31^. The study has illustrated the nature of the bacterial transfer from the early to mid-gut commensal bacteria to the pancreas. Yet none of these earlier studies have directly proved the presence of bacteria in the pancreatic cysts and its fluid. In the current study we have demonstrated that bacterial DNA is present in PCF. Although we do not have direct evidence that bacteria are alive in the PCF samples, the successful FISH analyses for some of the samples suggest that at least in part the bacterial population comprises of intact bacteria with undegraded 16S rRNA. PCF and the pancreatic duct borders represent therefore a niche that may become colonized by specific bacteria as demonstrated in the study.

The interactions between commensal and/or pathogenic bacteria and their metabolic products with hosts tissue can affect several diseases’ progression ^27,29,32^. Human serum derived antibody response against oral bacteria *Porphyromonas gingivalis* ATTC 53978 has been associated with an increased risk of pancreatic cancer by two folds ^33^. Furthermore, a study of Farell et.al., has showed that two oral bacteria, *Neisseria elongata* and *Streptococcus mitis,* can differentiate between pancreatic cancer and normal cases with 96.4% sensitivity and 82.1% specificity ^35^. In the same study, the oral bacteria *Granulicatella adiacens* and *Streptococcus mitis* differentiated between pancreatic cancer and pancreatitis with 85.7% sensitivity and 55.6% specificity, similarly in colorectal cancer the difference in bacterial Operational Taxonomic Units (OTU’s) were found between the normal, adenomatous and cancerous population, resulting in diagnosis of these three types in a person^34-36^. Analyses on the direct role of *Helicobacter pyroli* have shown that the bacterium was not associated with the development of pancreatic cancer ^37-40^. Two other studies, however, support indirect association between *H. pylori* and pancreatic cancer risk. *H. pylori* may represent high risk for the individuals with non-O blood types because *Helicobacter spp* DNA was found in 75% patients with pancreatic adenocarcinoma and in 57% of patients with neuroendocrine cancer, and in 60% of patients with chronic pancreatitis ^41,42^. Also the recent establishment of the role of *F. nucleatum, Bacteriodes spp.* in colorectal cancers in *in-vivo* and *Citrobacter rodentium,* in in-vitro conditions also shows the association of bacteria in general in cancer initiation and development ^26,33,36,43-46^ Microbiomes roles is being established in other cancers like urothelial *(Bacteriodes spp.)* ^47^ and prostate cancers *(Propionibacterium spp)* ^48,49^, liver cancer where the gut microbial metabolites were involved in the liver cancer progression ^50^.

Through this study we conclude, the bacterial presence was confirmed in the pancreatic cyst fluids, and as well-established via their copy number of 16S rRNA, Sangers sequencing and fluorescent in-situ hybridization. The kind of bacterial 16S rRNA shown, prove they were translocated from the mid and hind gut into the pancreas. Further analyses of the type of bacteria present in PCF can potentially help us to distinguish between various cyst types and the difference between the various grades of dysplasia.

It may also help in understanding the mechanism, whether it is a epiphenomenon or direct effect of how the presence of specific type of bacteria can potentiate the primary cystic lesions progressing towards pancreatic cancer. Furthermore, the data also raise a question of whether the bacterial population as seen in the pancreatic cyst fluid as well the cyst margins and duct margins could be a possible commensals of the pancreatic duct rather than bacterial invasion/translocation. A more comprehensive study is required for understanding this phenomenon.

## Acknowledgements

Hypothesis and experiment planning Sergey R konstantinov, experiment execution and manuscript written by Vilvapathy Narayanan, Manuscript reviewed by Wesley K Utomo, Sergey R Konstantinov, Marco J Bruno and Maikel P Peplenbosch. PCF fluid kindly provided by Gastroenterologists Henri Braat, J.W. Poley, A.D. Koch. FFPE blocks kindly provided by M. Doukas (Michael) pathology department, FFPE section were prepared by Juan Li Gastroentrolgy & Hepatology department, training and help in confocal microscopy provided by Gert-Jan Kremers and proteinase K was kindly provide by André Boonstra and Kim Kreefft Gastroentrolgy & Hepatology department

---

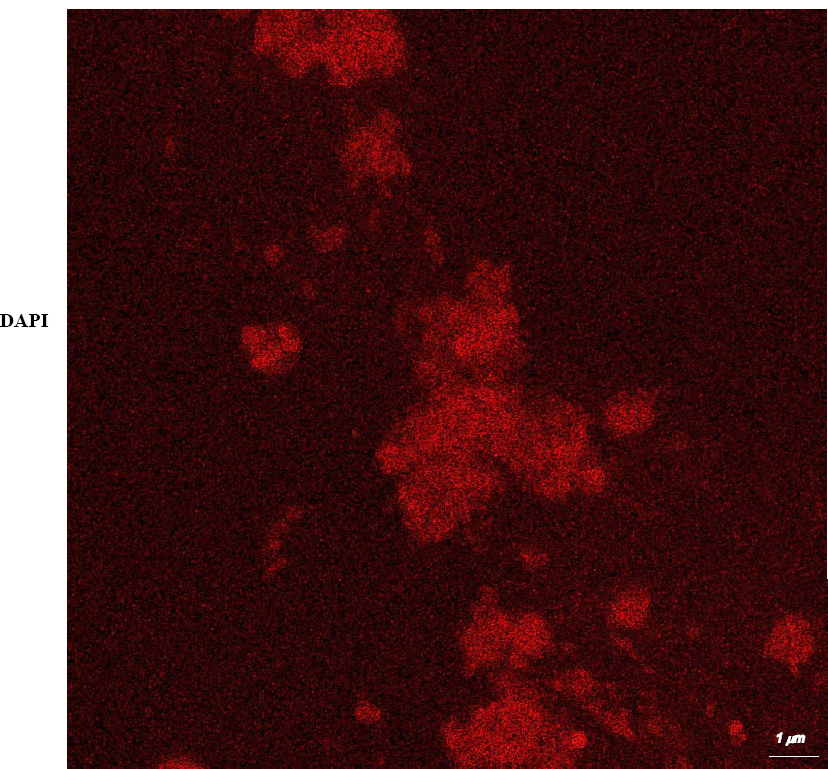
Bacterial 16S rRNA (EUB338 probe) in-situ hybridization in resected tissue.

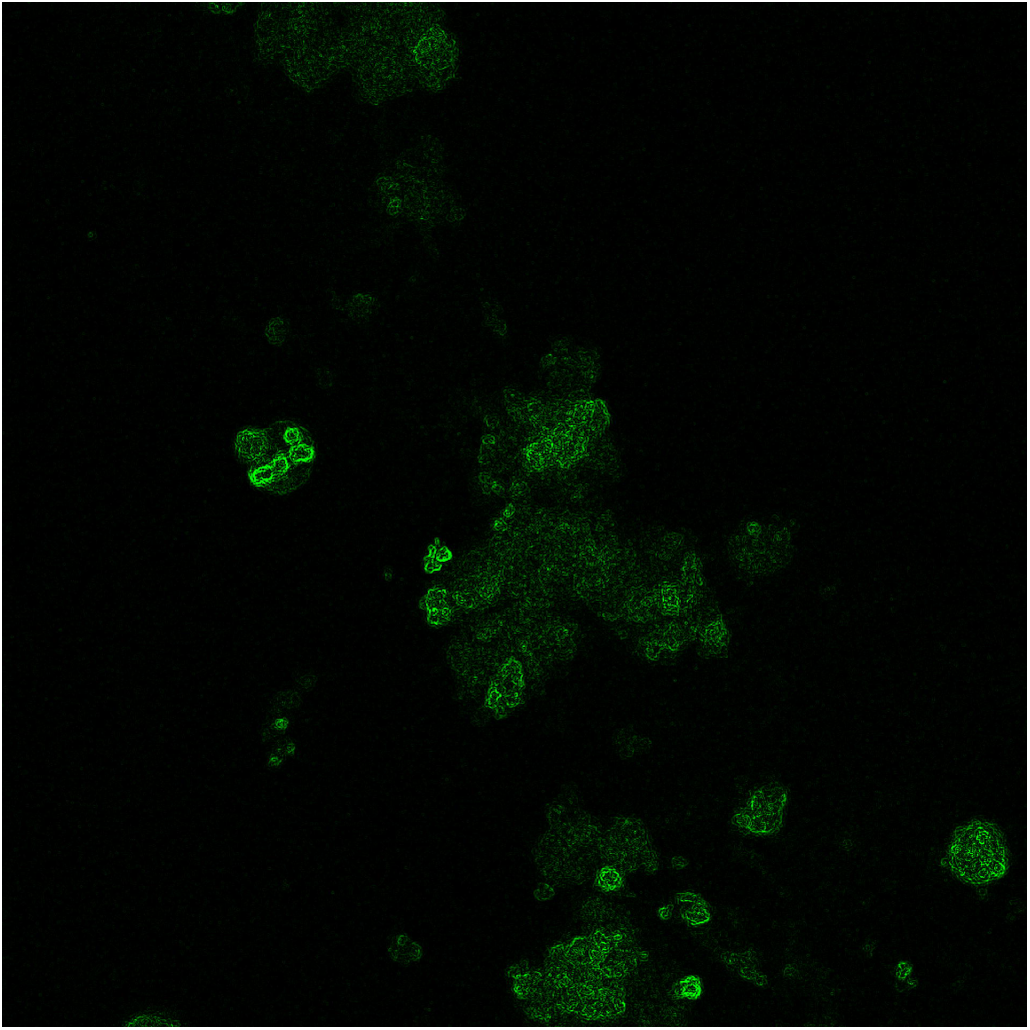

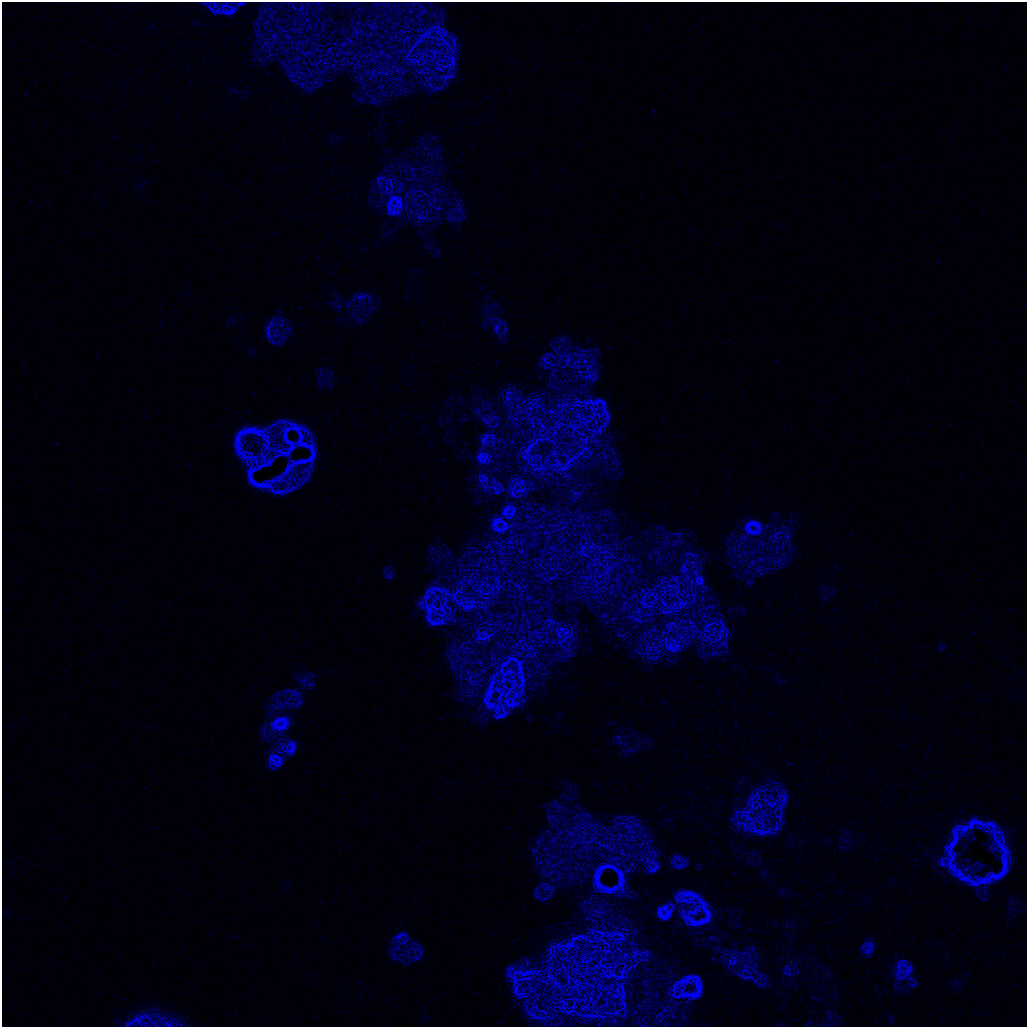

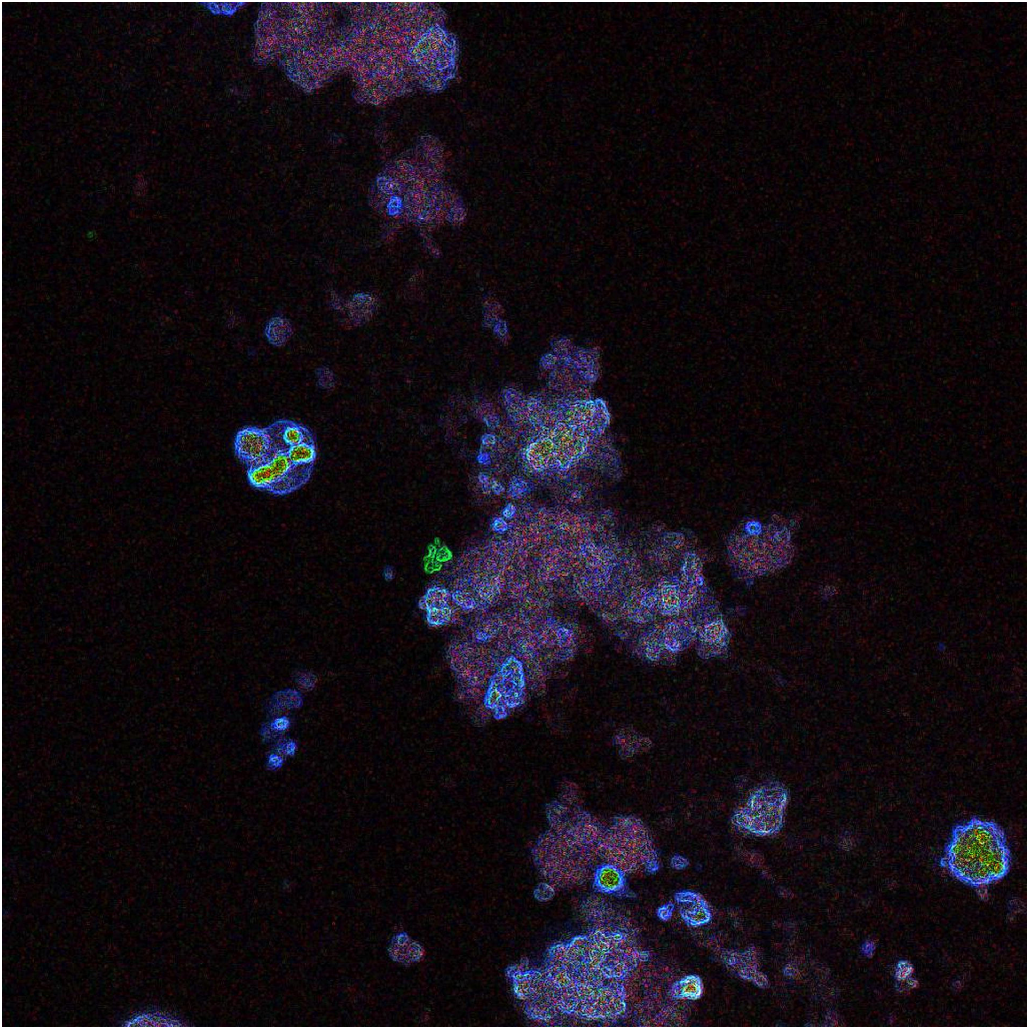

